# Molecular Adverse Outcome Pathways: towards the implementation of transcriptomics data in risk assessments

**DOI:** 10.1101/2023.03.02.530766

**Authors:** Marvin Martens, Anna Baya Meuleman, Jeroen Kearns, Chesley de Windt, Chris T. Evelo, Egon L. Willighagen

## Abstract

Adverse Outcome Pathways (AOPs) are designed to provide mechanistic insights into toxicological processes after exposure to a stressor and facilitate the replacement of animal studies with in vitro testing systems. Starting with a Molecular Initiating Event (MIE) and ending with an Adverse Outcome, the sequence of Key Events (KEs) spans across biological levels, but the majority of KEs in AOP-Wiki describe molecular and cellular processes.

That suggests that established transcriptome-wide studies can be used to validate or measure a multitude of KEs simultaneously and be a goldmine of useful information about cellular responses.

Currently, in toxicology, omics technologies are not widely applied in risk assessments of chemicals because of their complexity and lack of consensus on aspects such as standardization, analysis and interpretation. Given their value in hypothesis generation and ability to provide a vast amount of information about a biological response, the challenge lies in the acceptance of using transcriptomics data in regulatory risk assessments.

We here introduce molecular AOPs to define the connections between KEs of the AOP-Wiki and curated biological pathways in WikiPathways, thereby providing a new method for the analysis and interpretation of transcriptomics data to identify KE activation.

To study this, we performed case studies on liver steatosis and mitochondrial complex I inhibition, for which molecular AOPs were developed and public transcriptomics datasets were selected. Upon extension of the molecular AOP networks in Cytoscape, we mapped and analysed transcriptomics data, and calculated an enrichment score for individual KEs. Further interpretation of the data was done through the visualisation of the data on the specific molecular pathways.

Two molecular AOPs were developed and KEs were linked to the appropriate molecular pathways, allowing a detailed exploration of molecular processes with the selected transcriptomics datasets. This has shown us that we can verify the activation of specific MIEs and KEs, and assess progression across the AOP in the steatosis case study through variables in exposure time and dose.

These case studies have shown that transcriptomics data can be used for identifying the potential activation of KEs. However, it is also clear that extensive datasets are required to fully test the capabilities of molecular AOPs, and the process of linking molecular pathways and KEs can be challenging, not always allowing one-to-one mapping. While proven valuable to analyse and understand transcriptomic data, pathways linked to KEs appear to show inconsistent levels of activation and should be looked into and refined. More case studies are required to optimize the approaches used for the development and use of molecular AOPs with transcriptomics datasets.

## 1 Introduction

Adverse Outcome Pathways (AOPs) have become useful tools in risk assessments, as they provide a broad overview of events preceding a detrimental outcome. As such, they span multiple biological organizations, from the molecular level to the effects on a whole organism or even population. These types of pathways consist of three main concepts: Molecular Initiating Events (MIEs), Key Events (KEs), and Adverse Outcomes (AO). KEs are also linked with one another through Key Event Relationships (KERs), from molecular interactions to population dynamics [1]. The KEs in an AOP are not always sufficient on their own to lead to an AO. However, their strengths lie in that they are scientifically approved events that are essential to happen in the progression of the AOP. To be a well-defined AOP, the originating events from molecular interactions (i.e., MIE) should have a causal, measurable, and biologically plausible link towards an AO [2, 3]. With the evaluation of newly developed AOPs, one considers tailored Bradford-Hill criteria, in which the causality of observed association in epidemiological studies is determined [4]. This method considers biological plausibility, essentiality, and empirical support for these findings [4].

The practical aspect of these AOPs is that they are an integral part of risk assessment, where they act as scaffolds to collect and structure toxicological information at differing levels of the biological organizations, used to determine various apical AOs effectively after exposure to a stressor [1, 5]. This is also an essential part of Integrated Approaches to Testing and Assessment (IATA), which can include combinations of methods and integrating results of various types. AOPs can be used as a backbone to develop IATA [6].

However, current research is also focused on the quantification of AOPs in addition to the descriptive mechanisms, as risk assessors typically use numbers and thresholds to study the safety of substances and chemicals [7, 8, 9, 10]. These quantitative AOPs can be developed from qualitative AOPs, providing the quantitative descriptors or annotations for KEs and KERs, and can serve as predictors of adversity [7]. With the rise of genomic technologies and better biological system modelling, it is easier to hypothesize or develop new AOPs [11, 12, 13, 14].

The value of transcriptomics data in toxicological research is clear, and the technologies to produce the data are getting cheaper, faster and protocols better established [15, 16, 17]. However, transcriptomics technologies are not widely implemented in risk assessments [15]. This is due to the challenges in assessing and interpreting transcriptomics, and hurdles that exist in the general acceptance of transcriptomics data. For example, there is no consensus on the standardisation of the technologies and whether these can be validated. Furthermore, interpreting the data, distinguishing adaptive and adverse responses, investigating cause and consequence, and the amount of data required to make conclusions are challenges that remain to be solved [14, 12, 18].

For example, integrative approaches of publicly available data have been applied to develop AO networks of biological pathways and diseases which implement transcriptomics data [19], hypothesizing its support of evidence in risk assessments. Also, the integration of transcriptomics data has been studied for adverse pulmonary effects [13, 20]. The Organisation for Economic Co-operation and Development (OECD) has recently published a formal reporting framework to tackle the challenges of transcriptomics and metabolomics technologies in risk assessment applications. These provide guidance on the execution and reporting of data generation, processing, analysis, methodology and metadata [21].

It is expected that the combination of molecular pathways and AOPs allows the use of transcriptomics data for the measurement of KEs in AOPs and AOP Networks [12, 22]. Whereas biomarker genes and proteins exist for the validation of KEs, we believe that more elaborate molecular pathways can provide more robust evidence of KE activation using transcriptomics data [23, 24, 25]. Also, the connection between AOPs and molecular pathways offers additional insights into the molecular mechanisms underlying the more general KEs [26] and supports the biological plausibility of KERs.

One particularly well-studied AOP for applications of computational approaches and data integration is the AOP Network of liver steatosis that is caused through interactions with various nuclear receptors [27, 28, 29, 30]. The network is well-defined, and the receptor-specific MIEs allow the investigations of known agonists and antagonists of the receptors. This AOP network has been used to study a range of transcriptomics datasets of multiple chemicals known to activate specific MIEs [19, 31].

Another mechanism-unspecific AOP involves mitochondrial inhibition leading to AOs in multiple organs including the brain, liver, and kidneys [32]. A wide array of well-known chemicals are known to interact with the electron transport chain and disturb the production of ATP inside the mitochondria [33, 34]. While multiple downstream KEs are described for this, the most important KEs central in the AOPs are oxidative stress, unfolded protein response, and induction of cell death. This also counts for the well-established AOP of Parkinsonian motor deficits caused by mitochondrial complex I inhibitors [35, 36].

This paper provides a new method of applying transcriptomics data analyses and interpretations in the framework of AOPs. By doing so, the implementation of such data in risk assessment studies can be facilitated. To illustrate the value of this method, two case studies are performed, including the liver steatosis AOP Network and the AOP of mitochondrial complex I inhibition AOP leading to Parkinsonian motor deficits.

## 2. Methods

### 2.1Development of molecular AOP

To model molecular AOPs, PathVisio 3.3.0 [37] was used to draw and upload the molecular AOPs to WikiPathways [38], where they were tagged and stored in the AOP Portal (https://aop.wikipathways.org/, Figure 1). For the two case studies presented in this manuscript, the AOP-Wiki and relevant literature were used to construct AOPs. For each KE of the AOP, corresponding molecular pathways were identified in WikiPathways or, when necessary, developed based on the available scientific literature. The molecular AOPs exist as chains of Key Event nodes where each KE was annotated with the corresponding AOP-Wiki KE identifiers if available, linked with directed interactions representing KERs. Attached to the Event nodes are molecular pathway nodes containing identifiers of existing pathways in WikiPathways, using undirected interactions (see Figure 2).

**Figure 1:**
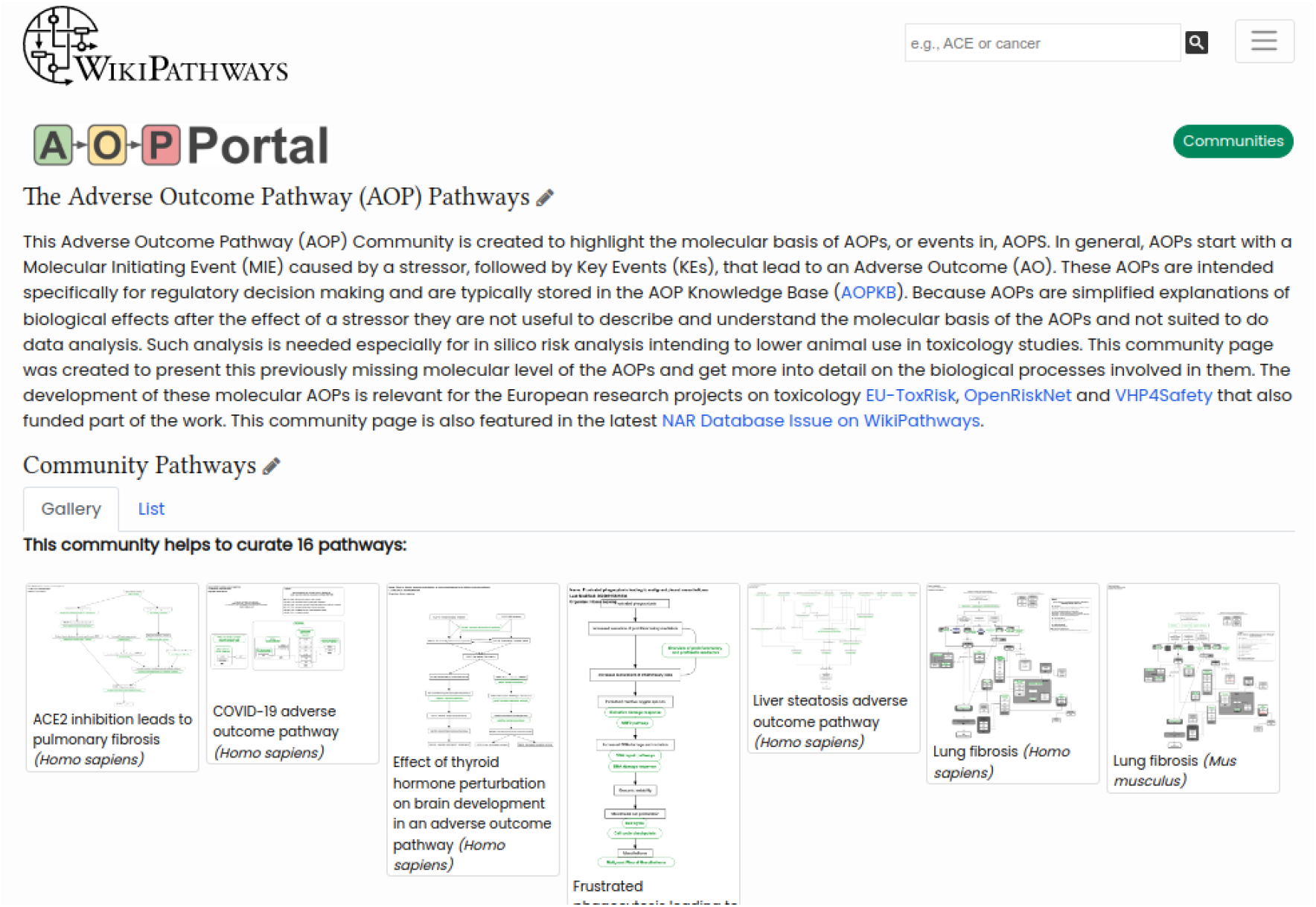
The AOP Portal on WikiPathways.org. On the portal, the AOP community can collaborate and comment on molecular AOPs and toxicity pathways.

**Figure 2:**
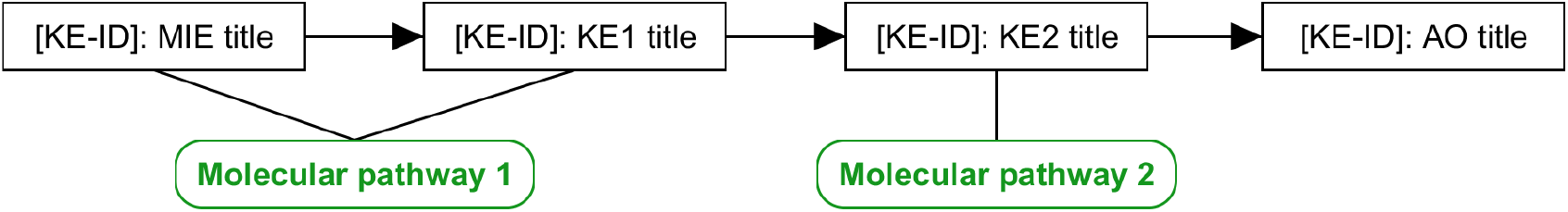
Conceptual illustration of a molecular AOP in WikiPathways. Black boxes are Key Events nodes, and green rounded nodes are Pathway nodes.

### 2.2 Datasets

To illustrate the implementation of gene expression data in molecular AOPs, publicly available datasets were selected in GEO [39] (see Table 1). Gene expression data of primary Human hepatocytes exposed to three Pregnane X Receptor (PXR) agonists from GEO dataset GSE90122 [40] was used in the case study of liver steatosis to explore and compare the effects of agonists of the PXR receptor, one of the MIEs of the liver steatosis AOP network. To explore the effect of time of exposure on gene expression in the liver steatosis case study, we also used data of HepaRG cells exposed to GW3965 (Liver X Receptor (LXR) agonist) from GEO dataset GSE123053 [41]. For a second case study of mitochondrial complex I inhibition, gene expression data from LUHMES cells (embryonic neuronal precursor cells) exposed to rotenone (mitochondrial complex I inhibitor) from the GEO dataset GSE116280 [42] was used to explore the effects of both time and dose on gene expression using molecular AOP. All datasets originated from GEO and were processed with GEO2R [43] to generate Log2 Fold Change (Log2FC) values and perform statistical tests to generate p-values for all reads in the comparison of samples with chemical exposure and without chemical exposure. Next, a custom Jupyter notebook was executed to calculate the average Log2FC values for each individual gene and Fisher’s combined probability test was used to calculate p-values for each gene.

**Table 1:**
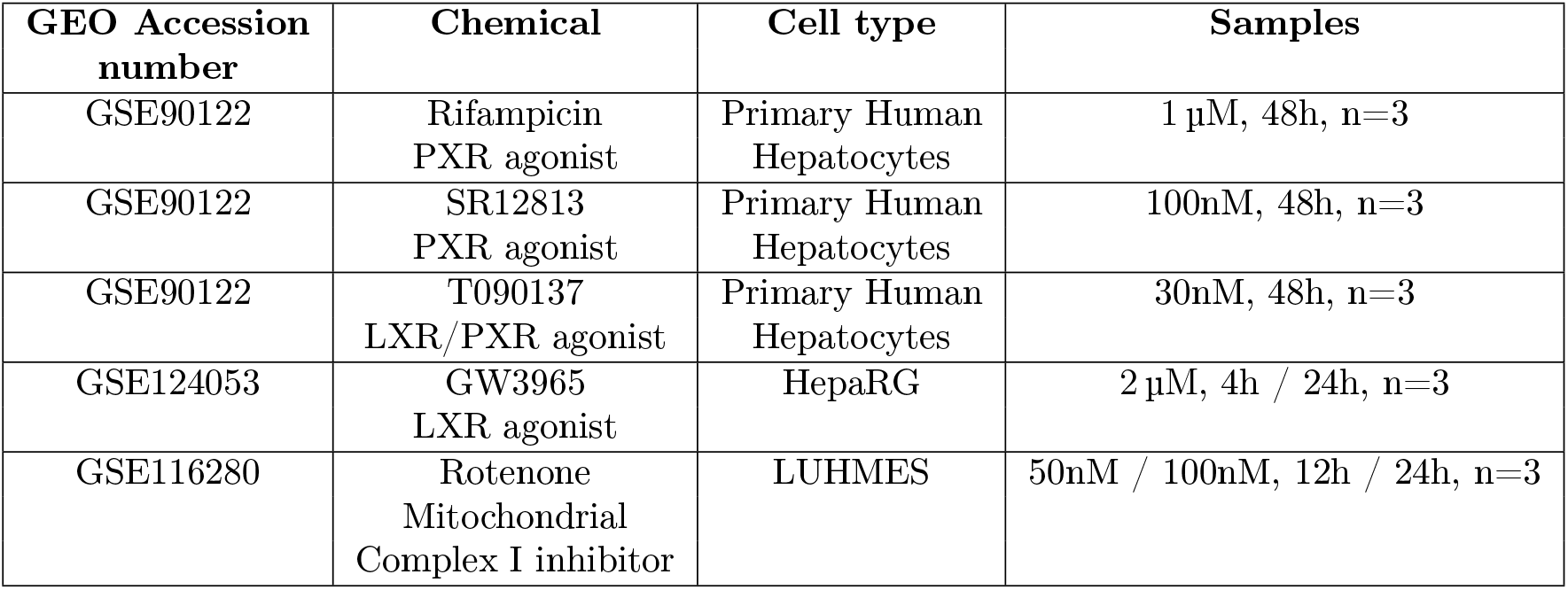
Overview of datasets used and detailed information on relevant samples.

| GEO Accession number | Chemical | Cell type | Samples |
| --- | --- | --- | --- |
| GSE90122 | Rifampicin<br>PXR agonist | Primary Human<br>Hepatocytes | 1 μM, 48h, n=3 |
| GSE90122 | SR12813<br>PXR agonist | Primary Human<br>Hepatocytes | 100nM, 48h, n=3 |
| GSE90122 | T090137<br>LXR/PXR agonist | Primary Human<br>Hepatocytes | 30nM, 48h, n=3 |
| GSE124053 | GW3965<br>LXR agonist | HepaRG | 2 μM, 4h / 24h, n=3 |
| GSE116280 | Rotenone<br>Mitochondrial<br>Complex I inhibitor | LUHMES | 50nM / 100nM, 12h / 24h, n=3 |

### 2.3 Cytoscape for AOP Network and data visualisation

In Cytoscape (version 3.8.2), the WikiPathways app (version 3.3.7) is used to import the molecular AOPs from WikiPathways as networks. Next, the network was extended with gene identifiers using the CyTargetLinker app (version 4.1.0) [44] with the WikiPathways Linkset (downloaded from https://cytargetlinker.github.io/pages/linksets/wikipathways, version 20210110. A custom visualisation was applied to distinguish Key Event nodes (orange diamonds) and pathway nodes (yellow squares).

Next, all datasets were imported into Cytoscape, and data were visualised on the gene nodes. These were colored with a blue-white-red gradient to represent the Log2FC values, where blue and red colors indicate the down- and upregulation respectively and white indicates no change. Node borders are highlighted in green for significantly altered expression levels (p-value < 0.05).

With the data visualised, the remaining genes without available data were removed. Also, in the case of the steatosis AOP network, the AOP was trimmed from irrelevant KEs which were not directly involved with the chemicals that were studied.

### Scoring Key Events

To quantify and assess the activation of KEs, the Enrichment Score (ES) is calculated based on the number of significantly affected genes present in molecular pathways linked to each of the KEs in the AOP, compared to the whole data set which was taken as the background. This is done using the formula

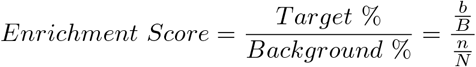

where:

*n* = number of differentially expressed genes in pathways linked to KE

*N* = total number of genes in pathways linked to KE

*b* = number of differentially expressed genes in the whole dataset

*B* = total number of genes in the whole dataset

Furthermore, a hyper-geometric p-value was calculated to indicate the statistical significance (p-value < 0.05) of the enriched pathways (ES > 1). The p-value was calculated using the formula

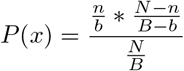

where:

*n* = number of differentially expressed genes in pathways linked to KE

*N* = total number of genes in pathways linked to KE

*b* = number of differentially expressed genes in the whole dataset

*B* = total number of genes in the whole dataset

### 2.5 Pathway data visualisation

For further exploration of the biological roles of differentially expressed genes, PathVisio was used to visualise the data on the molecular pathways linked to activated KEs. The same as for the molecular AOP in Cytoscape, the Log2FC values were visualised using a blue-white-red gradient in the left side of nodes where blue and red colors indicate the down- and upregulation respectively. Similarly, significance was visualised by a bright green color on the right side.

## 3 Results

In order to analyse the transcriptomic data in the case studies, molecular AOPs were developed for the AOP network of hepatic steatosis caused by multiple MIEs (wikipathways:WP4010 (identifiers.org/wikipathways:WP4010)) and the AOP of mitochondrial complex I inhibition leading to Parkinsonian motor deficits (wikipathways:WP4945 (identifiers.org/wikipathways:WP4945)).

### 3.1 Case study 1: liver steatosis

The molecular AOP network for liver steatosis consists of 30 KE nodes and 20 molecular pathways. In total, 578 unique genes were mapped to 20 of the KEs through the molecular pathways. The most highly connected gene is RXRA, being part of 8 of the molecular pathways. It codes for the Retinoid X Receptor, one of the nuclear receptor MIEs in the AOP network of liver steatosis.

For the datasets of PXR agonists, the PXR section of the liver steatosis AOP network was used for data visualisation and calculation of the KE enrichment scores (Figures 3, 4, and 5). In this subnetwork, 11 KEs were linked to 7 molecular pathways, containing unique 150 genes that were measured in the datasets of PXR agonists. The KE with the most significantly affected gene expression across the datasets of PXR agonists is the KE245: “PXR activation” and corresponding pathway wikipathways:WP2876, with thirteen out of 28 genes (48%) on average having significantly changed expression levels after exposure to one of the PXR agonists (Figure 6). This KE also has the highest enrichment scores of 12.83, 9.29, and 6.31 after exposure to Rifampicin, SR12813 and T090137, respectively (Table 2). The downstream KEs do not show consistency among the three PXR agonist data sets. The dataset of T090137 exposure notably shows a significant enrichment of most KEs except the on fatty acid lysis. Also of interest is the significant overexpression of the most connective (hub) genes of the network, including ACSL1, ACSL3, ACSL4, FASN, and ACACA, linked to at least three of the KEs in the network.

**Table 2:** Enrichment Scores for KEs by PXR agonists. Significance is indicated with an asterisk.

| Key Event | Pathway | Rifampicin | SR12813 | T090137 |
| --- | --- | --- | --- | --- |
| KE245: PXR activation | WP2876: Pregnane X Receptor pathway | 12.83* | 9.29* | 6.31* |
| KE471: FoxA2 inhibition | WP5066: Foxa2 Pathway | 4.79* | 2.89 | 3.31* |
| KE474: HMGCS2 Down Regulation | WP4718: Cholesterol metabolism | 0.53 | 3.85* | 2.94* |
| KE860: Decreased Mitochondrial Fatty Acid Beta Oxidation | WP368: Mitochondrial LC-Fatty Acid Beta-Oxidation | - | 3.40 | 3.25* |
| KE115: Increased FA Influx | WP5061: Fatty acid transporters | - | - | 3.07* |
| KE89: De Novo FA Synthesis | WP357: Fatty Acid Biosynthesis | - | 2.63 | 5.01* |
| Fatty Acid Lysis | WP3965: Lipid Metabolism Pathway | 0.83 | 1.00 | 1.90 |

**Figure 3:**
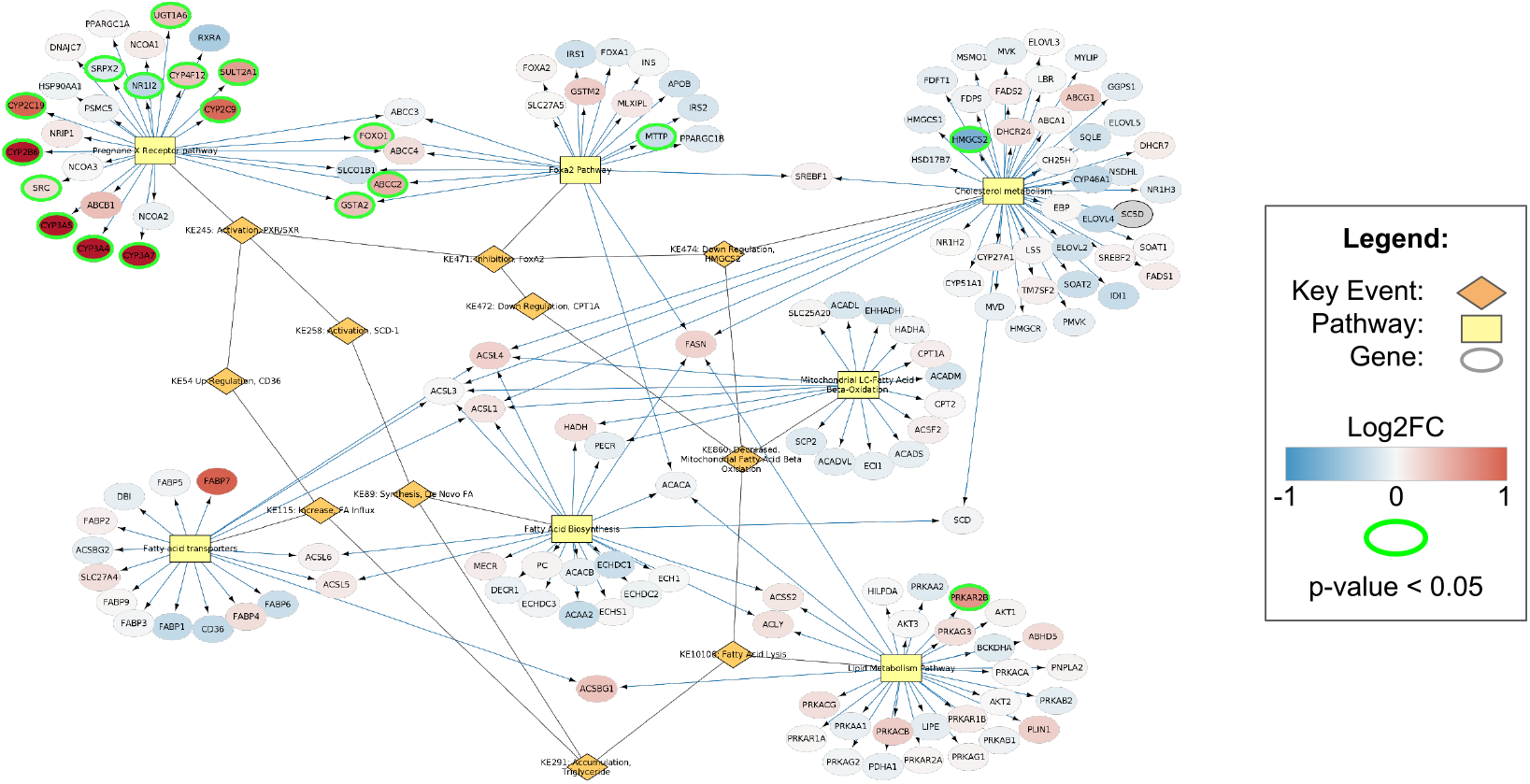
Gene expression data after Rifampicin exposure visualised on PXR AOP. Red and blue indicate up- and downregulation of gene expression (Log2FC), respectively. Green borders indicate significance (p-value < 0.05).

**Figure 4:**
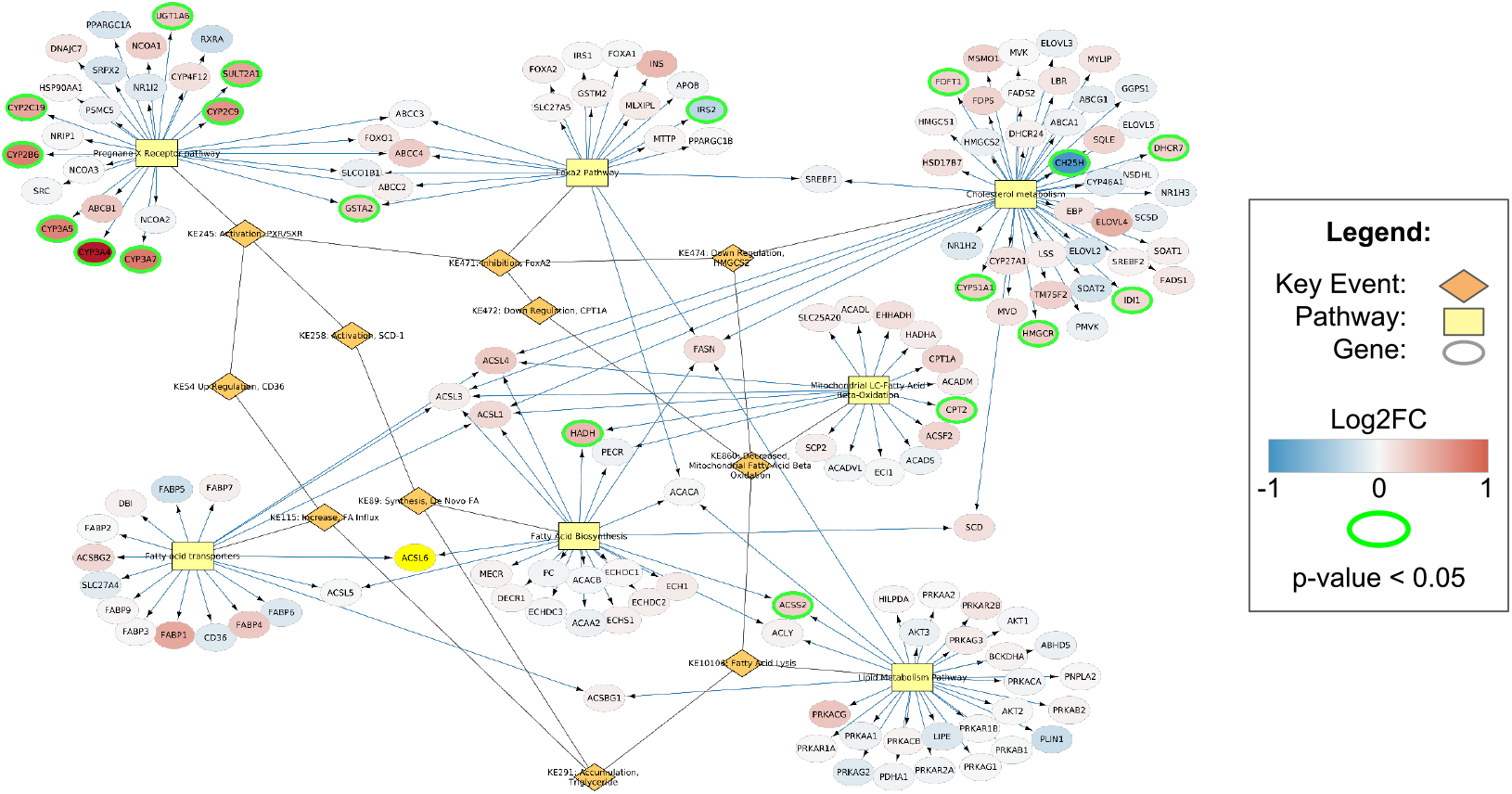
Gene expression data after SR12813 exposure visualised on PXR AOP. Red and blue indicate up- and downregulation of gene expression (Log2FC), respectively. Green borders indicate significance (p-value < 0.05).

**Figure 5:**
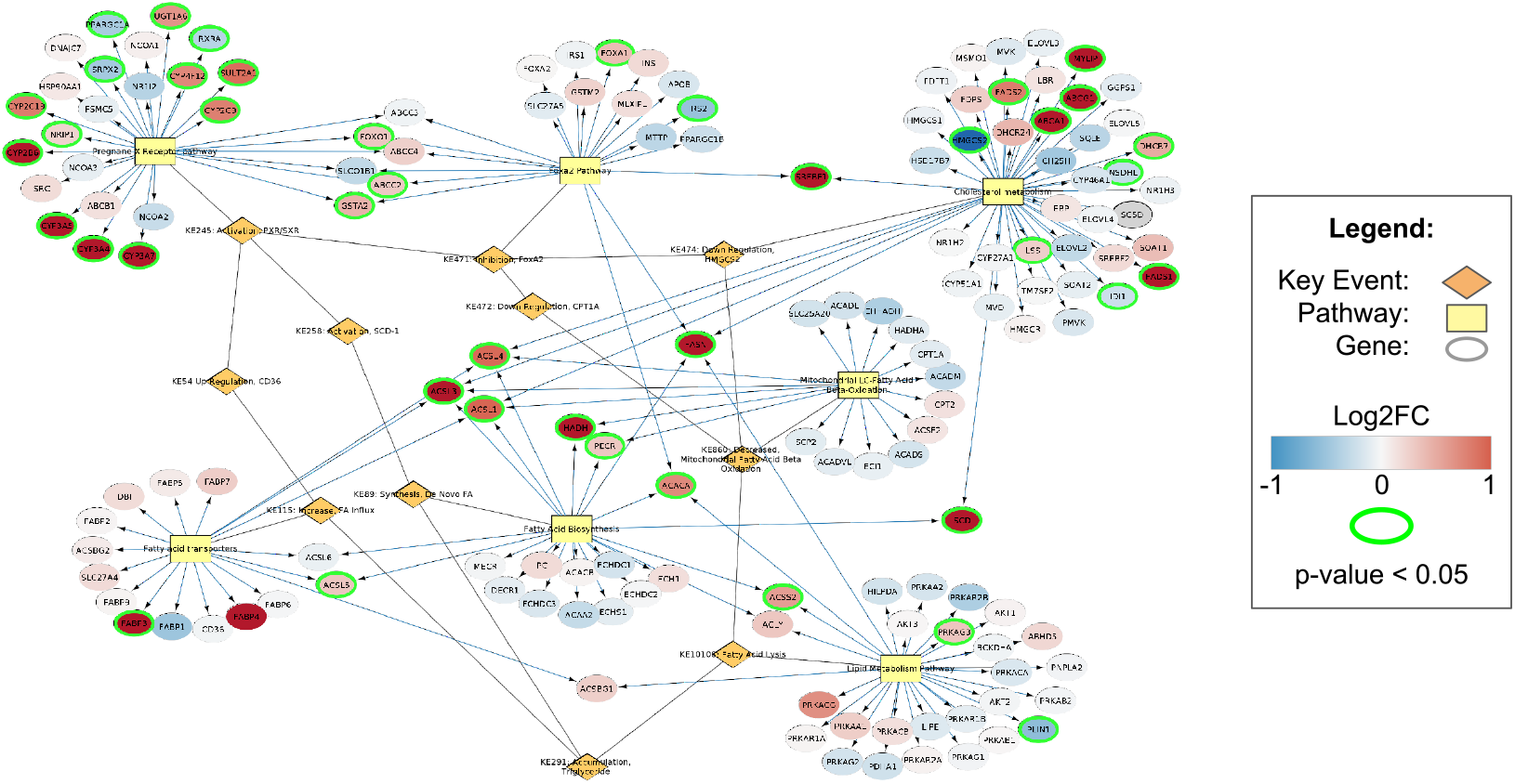
Gene expression data after T090137 exposure visualised on PXR AOP. Red and blue indicate up- and downregulation of gene expression (Log2FC), respectively. Green borders indicate significance (p-value < 0.05).

**Figure 6:**
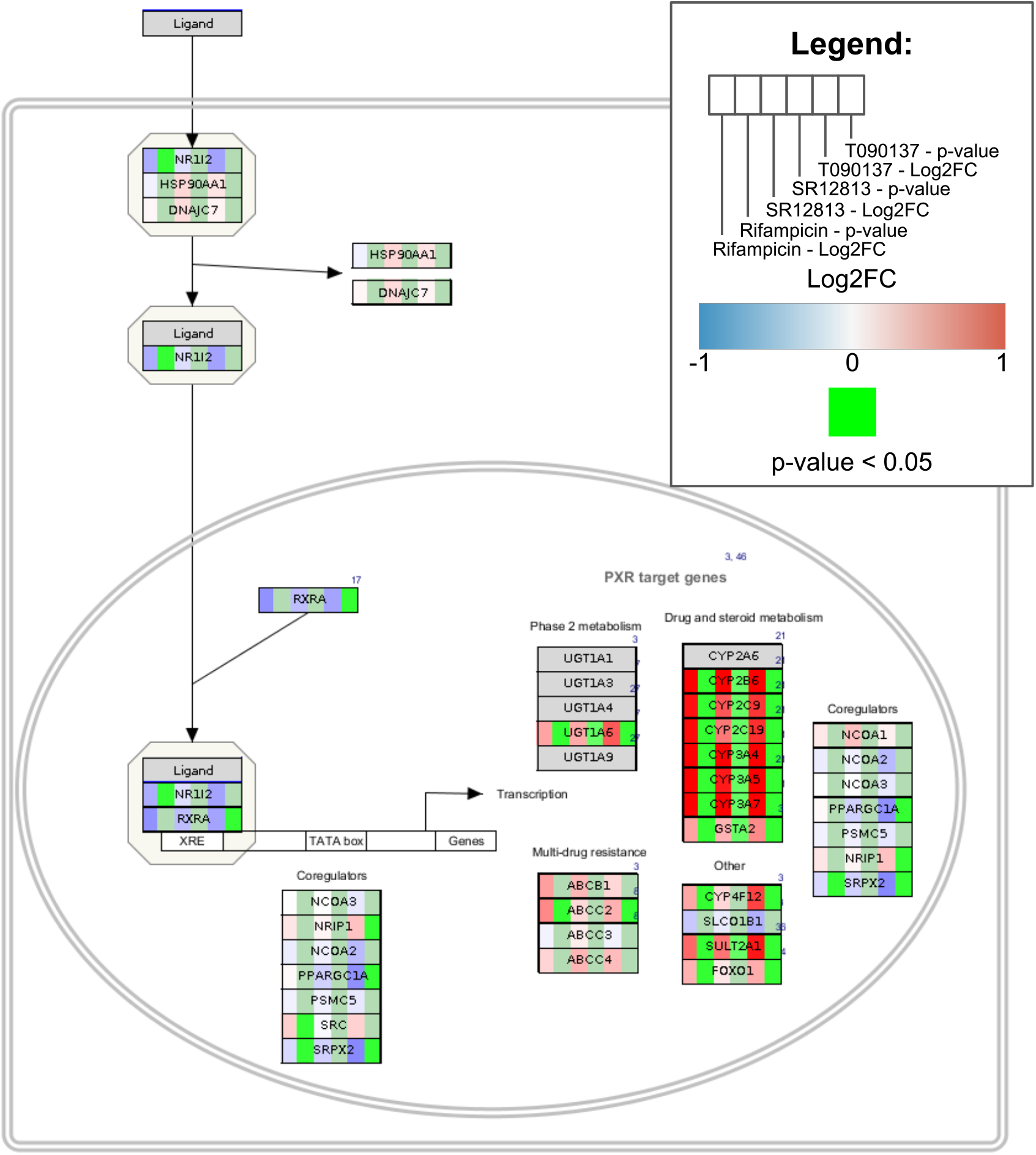
Gene expression data visualised on wikipathways:WP2876 (Pregnane X receptor pathway) for the three PXR agonists. From left to right, data is visualised for Rifampicin, SR12813 and T090137, in sets of Log2FC and p-values. Red and blue indicate the up- and downregulation of gene expression (Log2FC). Bright green marks indicate significance (p-value < 0.05).

Next to the PXR agonists, the second dataset was used to explore the effects of the LXR agonist GW3965 at different time points on the liver steatosis AOP. The AOP network was trimmed to have the MIE of LXR activation and all downstream KEs, leading to a new AOP network of six KEs, of which four were linked to five molecular pathways. In total, 226 unique genes were part of the network and were measured in the dataset. The most connected gene was FASN, which Molecular Adverse Outcome Pathways is involved in all five pathways, followed by SCD and SREBF1 which were part of four of the molecular pathways (Figures 7 and 8). A major difference between the time points is the number of significantly altered genes across the AOP, where nine genes are differentially expressed at four hours of exposure, and 27 genes at 24 hours of exposure. At four hours, the majority of the gene expression changes happen to the highly connected genes.

**Figure 7:**
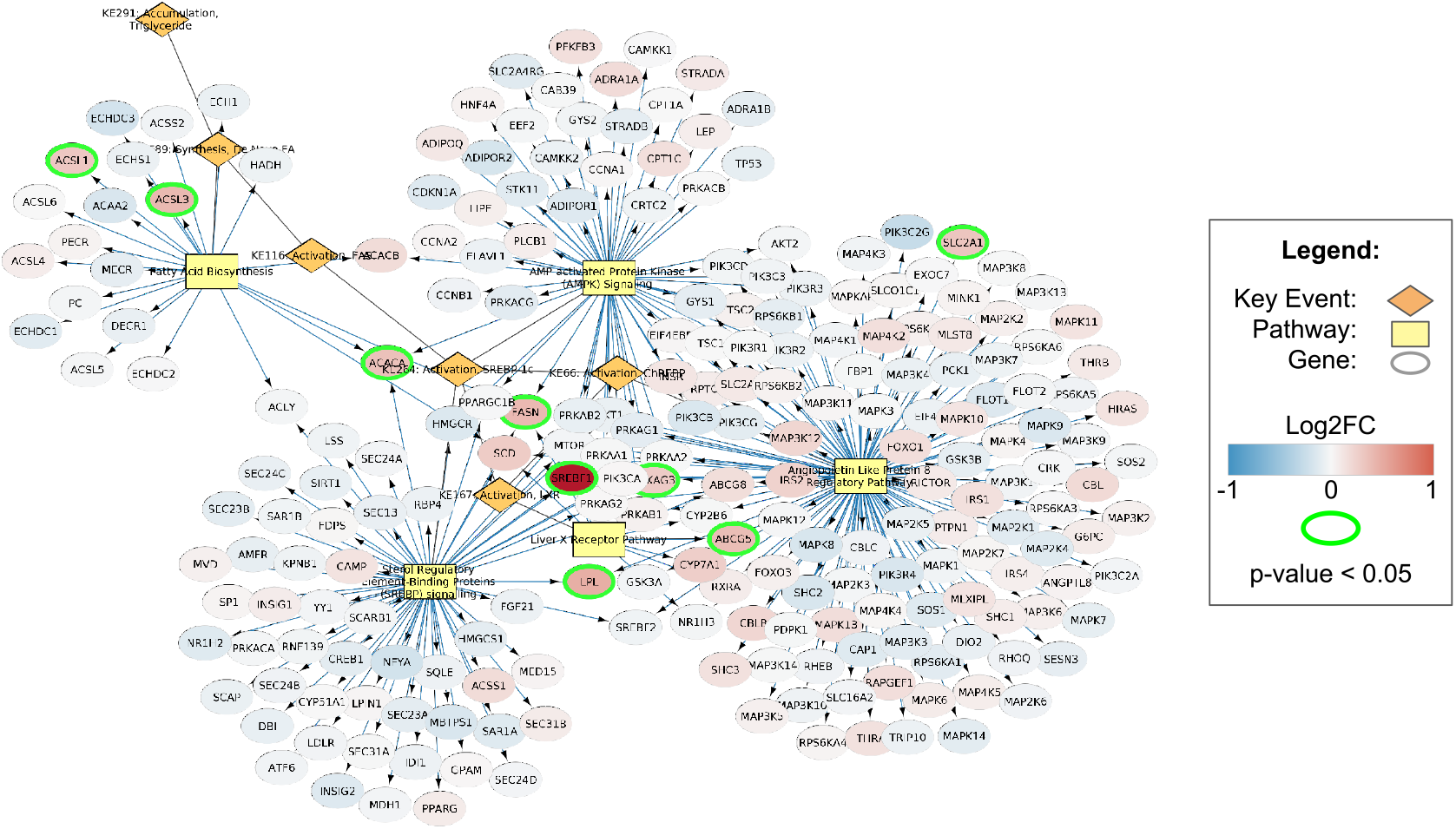
Gene expression data after 4 hour exposure to GW3965 (LXR agonist) visualised on LXR AOP. Red and blue indicate up- and downregulation of gene expression (Log2FC), respectively. Green borders indicate significance (p-value < 0.05).

**Figure 8:**
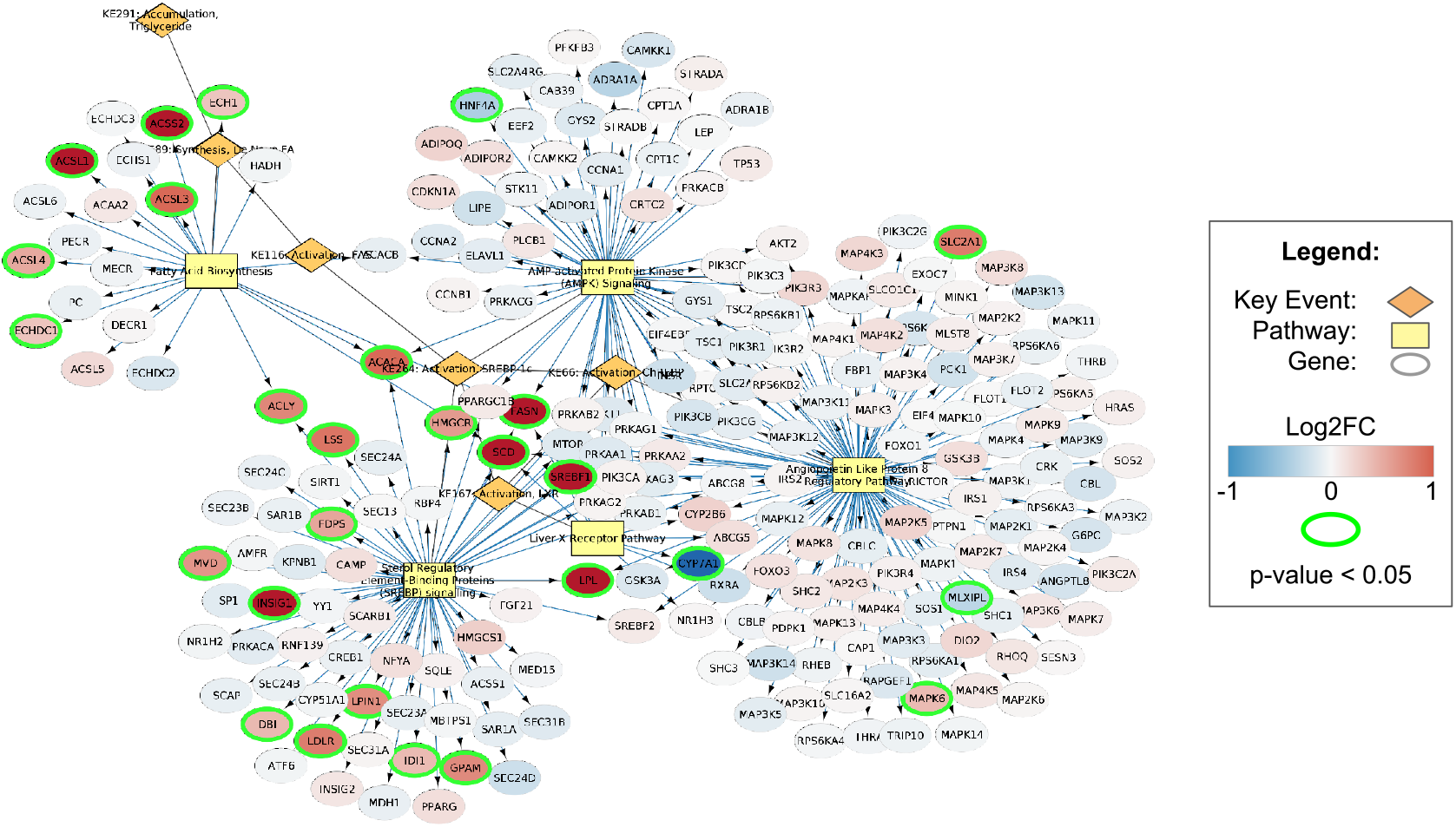
Gene expression data after 24 hour exposure to GW3965 (LXR agonist) visualised on LXR AOP. Red and blue indicate up- and downregulation of gene expression (Log2FC), respectively. Green borders indicate significance (p-value < 0.05).

From the data on the LXR AOP, it can be noted that the MIE of LXR activation and the KE of Fatty Acid Biosynthesis were significantly enriched at both time points (Table 3). However, the largest KE of SREBP-1c activation, linking to 113 genes, is only significantly enriched at 24h of exposure to GW3965. This difference is most visible in the SREBP pathway, whereas the AMPK pathway does not differ between the time points.

**Table 3:** Enrichment Scores of KEs after exposure to GW3965 (LXR agonist). Significance is indicated with an asterisk

| Key Event | Pathway(s) | GW3965 4h | GW3965 24h |
| --- | --- | --- | --- |
| KE167: LXR activation | WP2874: Liver X Receptor Pathway | 12.31* | 11.46* |
| KE66: ChREBP activation | WP3915: Angiopoietin Like Protein 8 Regulatory Pathway | 1.72 | 1.60 |
| KE264: SREBP-1c activation | WP1982: SREBP signaling<br>WP1403: AMPK signaling | 1.63 | 3.88* |
| KE89: De Novo FA synthesis | WP357: Fatty Acid Biosynthesis | 6.71* | 11.72* |

### 3.2 Case study 2: mitochondrial inhibition

The second case study was focused on the AOP of mitochondrial complex I inhibition leading to Parkinsonian motor deficits. The developed molecular AOP contains seven KE nodes and seven pathway nodes. When extended using CyTargetLinker, a total of 199 gene nodes are added to the molecular AOP network and are measured in the datasets of Rotenone exposure to LUHMES cells. Of all the genes in this network, approximately 25% show significantly altered gene expression levels upon exposure to Rotenone.

The datasets for rotenone on LUHMES cells were visualised and they show the significantly altered gene expression for all KEs, where 50nM dose exhibits a stronger effect in the early KEs of mitochondrial complex I and oxidative phosphorylation, and the 100nM dose causes a higher number of gene expression changes in the AO of Parkinsonian motor deficits (Figures 10 and 11). The only KEs that are significantly enriched are the binding and inhibiting of mitochondrial complex I at 50nM Rotenone exposure (Table 4).

**Table 4:**
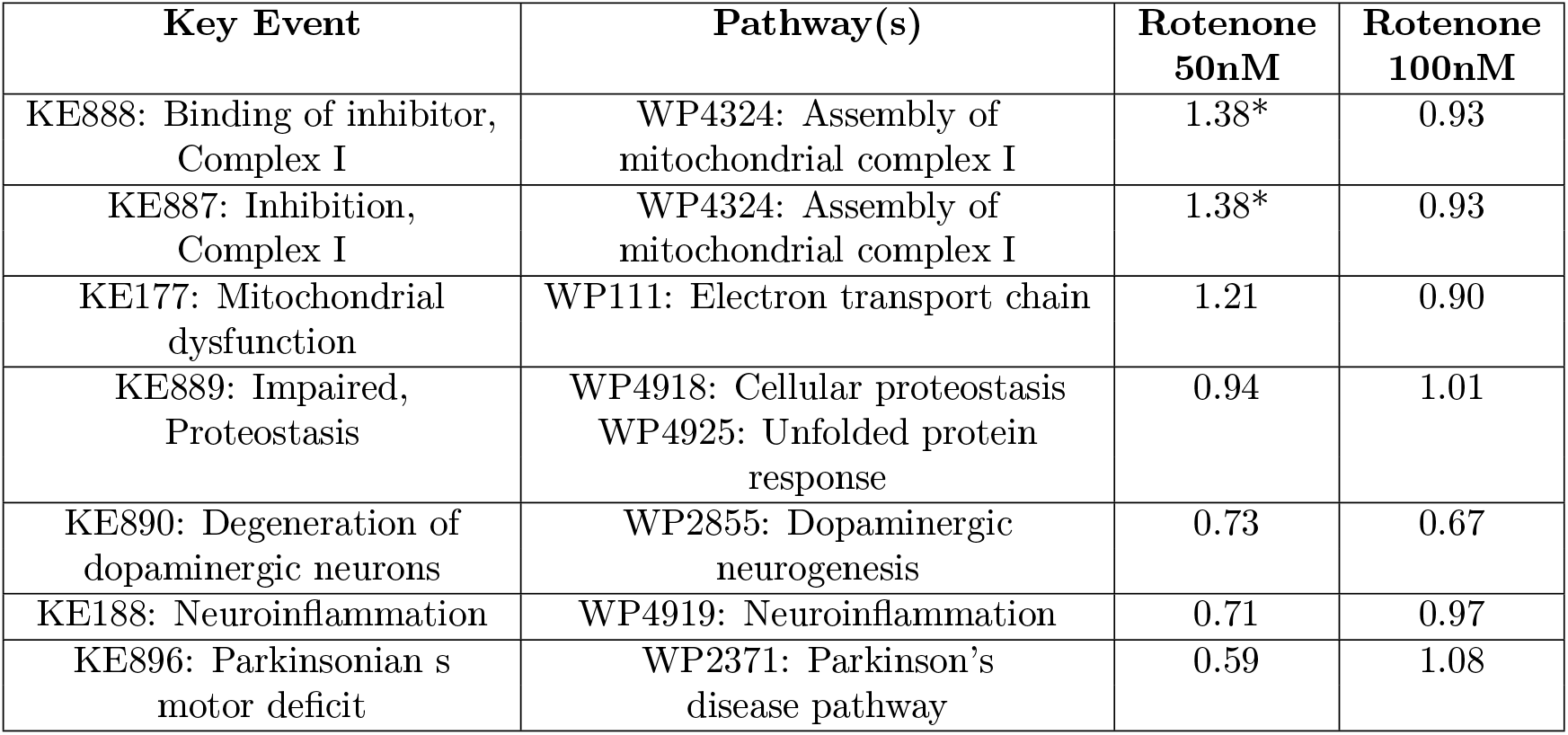
Enrichment Scores for KEs by exposure to Rotenone at 50nM and 100nM. Significance is indicated with an asterisk

| Key Event | Pathway(s) | Rotenone 50nM | Rotenone 100nM |
| --- | --- | --- | --- |
| KE888: Binding of inhibitor, Complex I | WP4324: Assembly of mitochondrial complex I | 1.38* | 0.93 |
| KE887: Inhibition, Complex I | WP4324: Assembly of mitochondrial complex I | 1.38* | 0.93 |
| KE177: Mitochondrial dysfunction | WP111: Electron transport chain | 1.21 | 0.90 |
| KE889: Impaired, Proteostasis | WP4918: Cellular proteostasis<br>WP4925: Unfolded protein response | 0.94 | 1.01 |
| KE890: Degeneration of dopaminergic neurons | WP2855: Dopaminergic neurogenesis | 0.73 | 0.67 |
| KE188: Neuroinflammation | WP4919: Neuroinflammation | 0.71 | 0.97 |
| KE896: Parkinsonian s motor deficit | WP2371: Parkinson's disease pathway | 0.59 | 1.08 |

**Figure 9:**
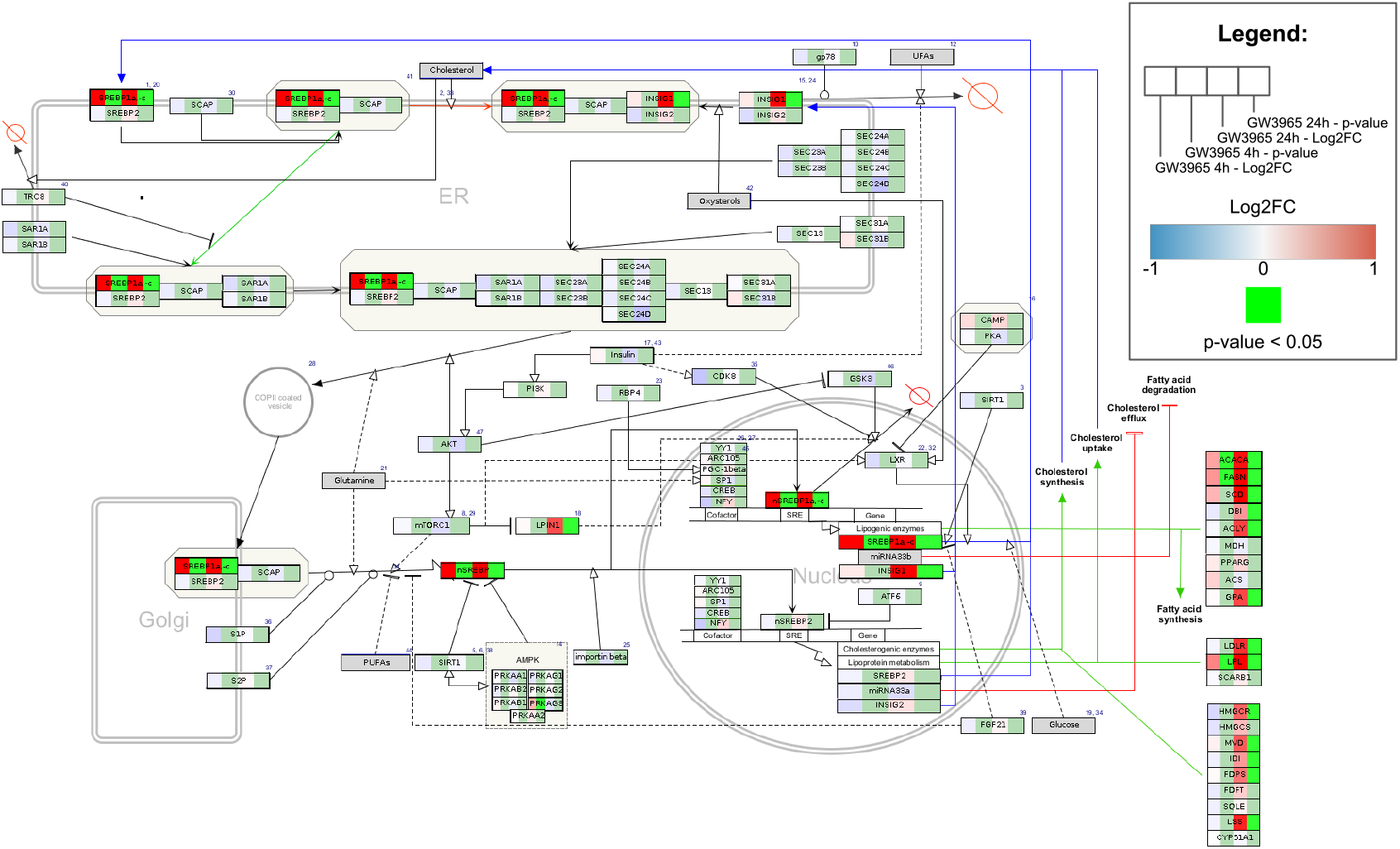
Gene expression data after exposure to GW3965 (LXR agonist) visualised on the SREBP signaling pathway (wikipathways:WP1982). The left side of each data node represents data from 4h exposure, and the right side represents the data from 24h exposure. Red and blue indicate the up- and downregulation of gene expression (Log2FC). Bright green marks indicate significance (p-value < 0.05).

**Figure 10:**
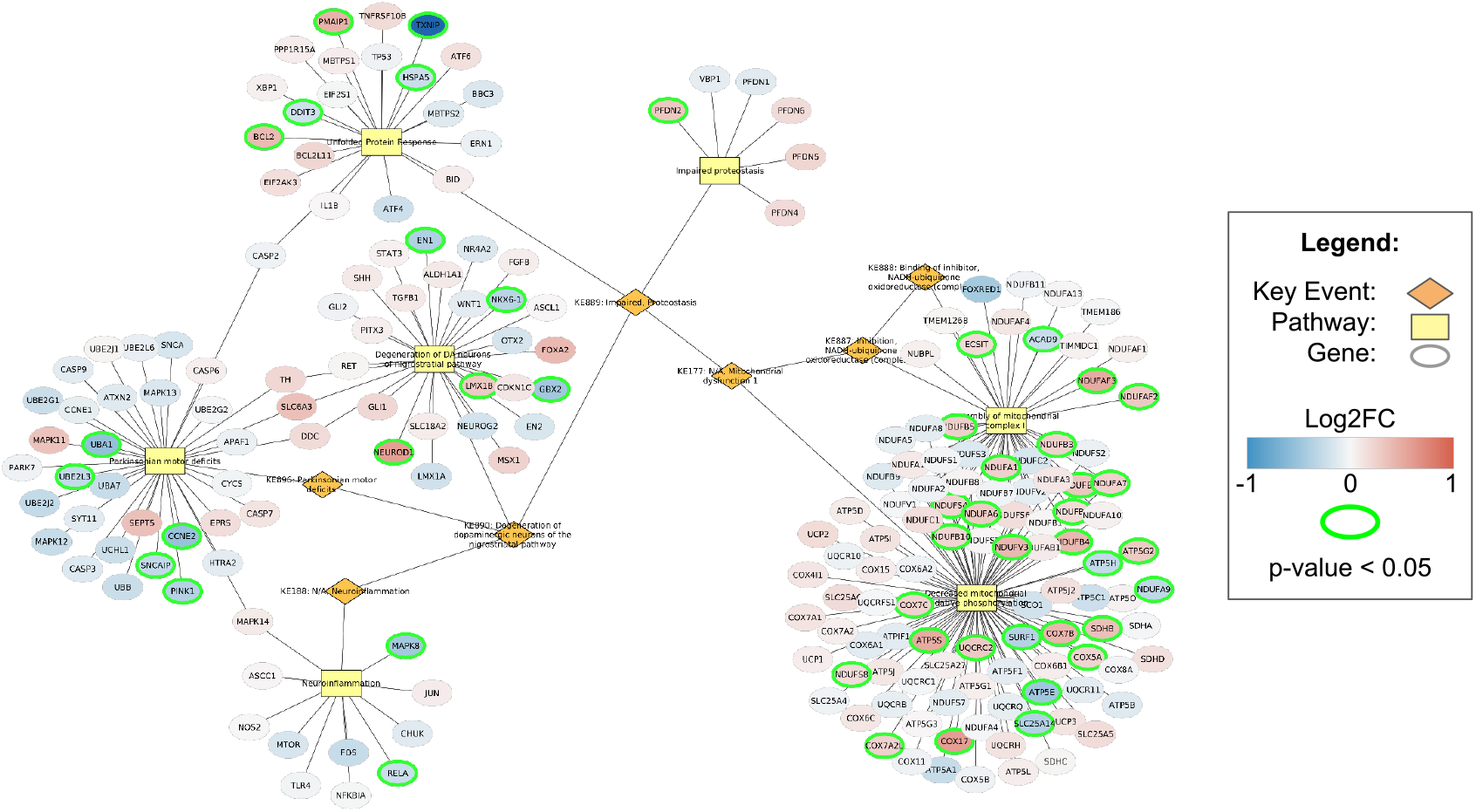
Rotenone (50nM) data visualised on mitochondrial complex I inhibition AOP. Red and blue indicate the up- and downregulation of gene expression (Log2FC). Bright green marks indicate significance (p-value < 0.05)

**Figure 11:**
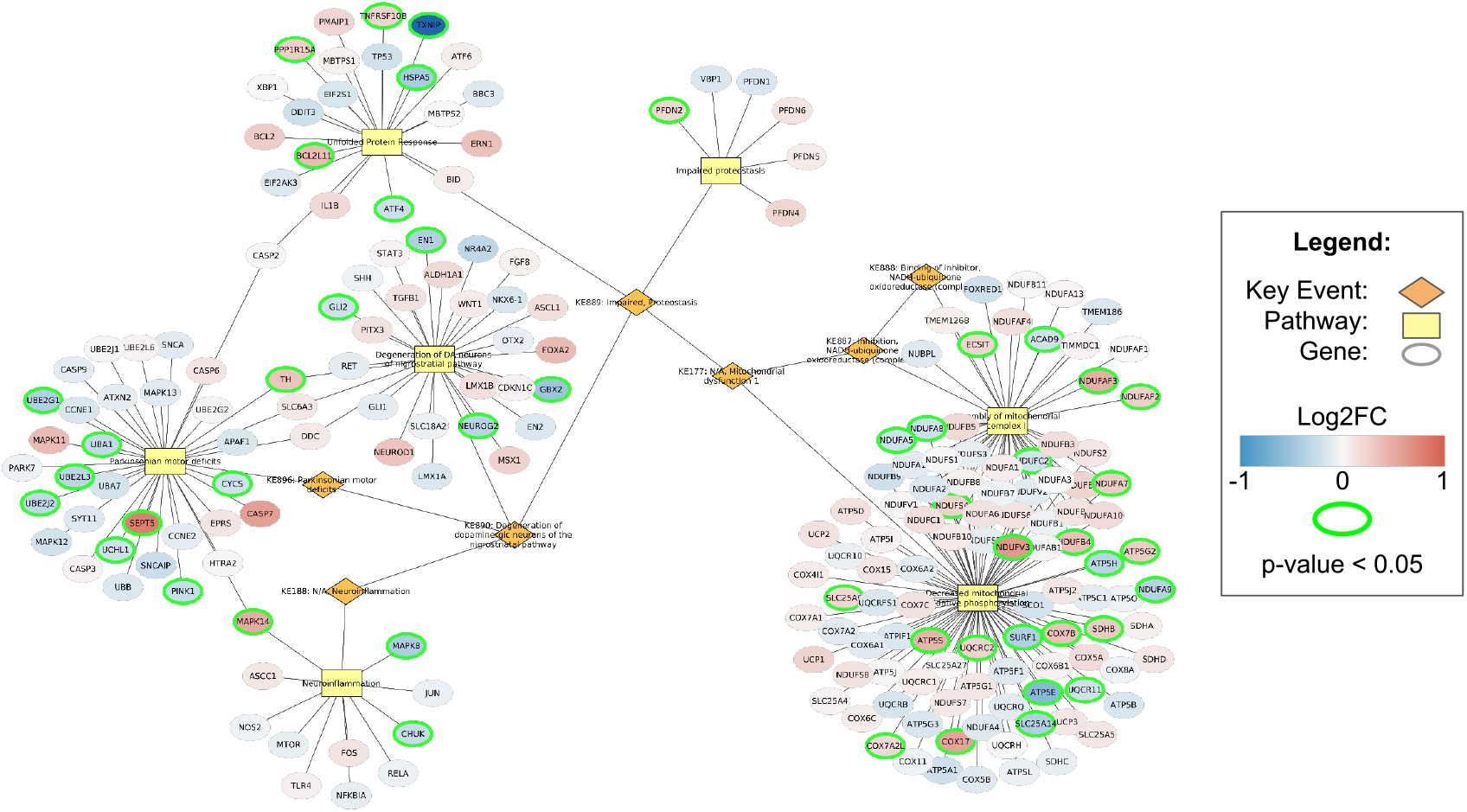
Rotenone (100nM) data visualised on mitochondrial complex I inhibition AOP. Red and blue indicate the up- and downregulation of gene expression (Log2FC). Bright green marks indicate significance (p-value < 0.05)

## 4 Discussion

With this work, we have created the molecular counterparts of AOPs, tightly linked to matching AOPs in the AOP-Wiki through the linking of molecular pathways explicitly to KEs. The addition of molecular pathways to KEs to analyse transcriptomic data allows a broader sense of the effects of toxicants when compared to biomarkers for process activation, which most often provide limited information and are used to measure a single KE. Therefore, these molecular AOPs can be used to get a more thorough understanding of traditional AOPs by exploring molecular pathway models, but they also provide a method of visualising and analysing omics datasets, which was explored in this manuscript.

When looking at gene expression changes upon exposure to a toxicant, the predictive value of each individual gene in the network could differ. This is where pathway models can provide additional insights into the overall connectivity of genes and their roles in pathways [45, 46].

The first case study was focused on the liver steatosis AOP, a widely studied and well-established AOP [27, 47, 28, 48], starting with a range of nuclear receptors known to be involved in maintaining lipid balance in the liver, including LXR, PXR, PPAR, and CAR, among others [48]. With the case study, we showed that molecular AOPs can be used to visualise transcriptomic datasets and perform enrichment analyses of KEs. Based on the results, we can clearly identify the activated MIEs and make comparisons between chemicals or exposure scenarios. For example, the dataset of GW3965 could be used to investigate the differences in gene expressions at two time points. By performing an enrichment analysis, many processes can be assessed simultaneously and generate hypotheses of KE activation.

Although the molecular AOP can highlight which KEs are affected based on pathway-level gene expression changes, the mapping of KEs to molecular pathways does not always fill its purpose. Some of the KEs in the liver steatosis AOP network involve single gene expression changes, rather than processes being affected, and for those KEs a single gene expression biomarker might already be sufficient. For example, the expression level of HMGCS2, which plays an essential role in cholesterol metabolism and ketogenesis [49], is significantly decreased after exposure to T090137 and Rifampicin. However, since the focus of our analysis lies on the molecular pathways to expand our biological understanding, the KE is only regarded as significantly enriched within the dataset of T090137 exposure. This can be a limitation for KEs that have a clear transcriptional marker gene or gene set, or KEs that are confined to single gene expression changes rather than processes. This distinction of single markers and pathways would also be important to address for the KE of SCD-1 (SCD) activation, which plays a role in the regulation of energy metabolism and lipid synthesis, and has significantly increased expression levels after exposure to T090137. In the molecular AOP network, however, it is linked to the Cholesterol metabolism and Fatty Acid Biosynthesis pathways, and not the specific KE of SCD-1 activation. The data shows that only T090137 exposure led to the enrichment of the majority of KEs whereas the other PXR agonists only led to the enrichment of a handful of KEs. This can be due to the dual agonistic role of T090137, as it is also an LXR agonist [50], affecting the downstream KEs through multiple pathways.

On the other hand, the KEs of CD36 upregulation, being a fatty acid transporter [51], does not show up as significant in the gene expression data but it is part of the Fatty Acid Transporters pathway which is significantly enriched after T090137 exposure. This is also the case for the KE of CPT1A downregulation which does not show up in the data of the PXR agonists. However, it is part of the pathway linked to the downstream KE of Decreased Mitochondrial Fatty Acid Beta Oxidation, which is significantly enriched after T090137 exposure.

As discussed, there can be value in regarding single transcriptional marker genes to investigate KE activation. Since individual or groups of (computed) transcriptional biomarkers can provide great insights into the activation of individual processes [52], further developments of this approach should focus on combining the molecular AOPs with stress response marker genes relevant to individual KEs. The value of using biomarkers or defined gene sets to explore the effects of toxicants and xenobiotics has been shown in various studies related to AOPs and KE activations. For example, well-described transcription factor modulations can be explored with predictive gene sets [23] or well-established transcriptional biomarkers based on responsive transcription factors [53]. An approach to combine the well-studied and carefully selected expression biomarkers for KE activation and molecular pathways can be most informative to validate the activation of KEs and understand the biological responses at the cellular level in more detail.

With the case studies presented in this work, we limited the investigation to AOPs that involve only a single cell type. While that can be the case for sequential KEs that are mostly on the molecular, cellular or tissue level, KEs can also involve cellular communication, such as the secretion of signalling molecules or recruitment of inflammatory cells, as inflammation plays an important role in toxic adverse outcomes [54]. For such AOPs to be fit for use as molecular AOP, multiple datasets would be required to assess the KEs across cell types, making the overall assessment more complex. However, the flexible nature of WikiPathways, the identifier handling by BridgeDb and the integration of these tools in Cytoscape facilitate the integration of data and the model.

Based on our analyses and calculations of enrichment scores, the case studies provide a great insight into the usability and value of transcriptomics data within the molecular AOPs, showing the potential activations of KEs. With the visualisation and enrichment calculations, this method provides a quick overview of the overall activation of pathways that are linked to the KEs of interest. Also, it shows the interplay between processes within the AOP, highlighting central, highly connected genes, whose role can be further explored in the molecular pathways in which they exist. This is for example clearly visible in the PXR AOP, with various members of the ACSL family being involved in multiple KEs, as well as FASN being the most connected gene in the network. A significant alteration of their expression levels is expected to have a stronger impact on the overall assessment within the AOP when compared to genes that are part of only a single pathway or KE.

With the case study on liver steatosis, we focused on highly specific MIEs with well-studied stressors, and these show the activation of MIE-linked pathways based on gene expression data, which was consistent across the chemicals that we investigated for the different MIEs. Furthermore, with the multiple time points in the dataset for stressor GW3965, it is clear that the exposed cells progress through the KEs of the AOP towards the AO, which is promising for the application of time series exposure data on molecular AOPs. However, more extensive datasets with additional time points and doses would be required to assess the progression through the AOP based on gene expression data.

The case study of mitochondrial complex I inhibition by rotenone is showing the challenges of interpreting transcriptomics data and variations in dose-response data. While in low-dose exposure we see an abundant upregulation of genes involved at the early KEs to counter the initial inhibition of mitochondrial complex I, this is not as clear in the higher-dose exposure. This suggests a switch in response from recovery towards adverse, but this is not clearly visible in the late KEs which represent stress response processes, none of which have been affected significantly. With the current calculation of KE enrichment, the whole dataset is taken as the background data, of which approximately 25% had significantly altered gene expression levels. Since the datasets of the liver steatosis case study contain between 3% and 9% of significantly affected genes, it could be that the disturbance of cellular energy production causes many more processes to be affected. This relates to our approach to developing the molecular AOPs because we limit ourselves to the known KEs and do not include additional processes or responsive (feedback) pathways in the molecular AOP model. This constitutes a challenge in our approach, in which the pathway’s level of detail and focus originate from their initial creation by the many contributors [55]. Since the method for creating the molecular AOP involves the inclusion of existing pathways developed by the community, curation might be necessary to ensure the expected quality of pathways to support KEs and to explore the outputs of the analyses.

Another challenge lies in the distinction between causative and responsive processes to the toxicity of stressors. For example, the MIEs of the steatosis AOP are nuclear receptor activation pathways that cause downstream effects upon their activation or inhibition, and clearly show transcriptional changes based on the exposure to stressors. However, the effects on pathways linked to downstream KEs are much more subtle and respond to the changes that occur because of the activated MIE. On the other hand, the case study of mitochondrial inhibition does not provide a clear-cut activation of a pathway related to the MIE but instead consists mostly of responsive pathways to adapt to the new situation caused by the stressor. To develop useful molecular AOPs, one needs to be aware of the expected effects on the transcriptional level, which does not always match the KE description in the AOP-Wiki, where feedback loops and context are less present.

Whereas some KEs describe the activation of cellular responses and processes and are therefore easily linked to their molecular pathways in WikiPathways, other KEs can be more simple or more complex. For example, KEs can merely describe individual molecular interactions such as receptor activation, or describe the larger, more general interplay of processes, such as cell death where the measurement is focused on cellular viability. This varying level of complexity should also be represented in molecular AOPs.

Based on these results, taking into account the limitations, we find that we can explain the biological plausibility of KE activation by visualizing experimental transcriptomics data to the molecular pathways underlying the KEs. The extension of AOPs with molecular markers, gene sets, or pathways would be an essential step towards the integration of high-throughput transcriptomics data into risk assessment studies. With these technologies getting cheaper, faster, and more reliable, they are becoming more frequently used in toxicological research. Hence, a framework to connect the established AOPs on AOP-Wiki with molecular entities through biological pathways is the logical intermediate.

Future work should focus on expanding the molecular AOP contents and on validating their utility through additional case studies and comparing the outcomes to other methods to measure KE activation using gene expression data, such as the TXGMAPr tool [56] or AOP fingerprints [57]. When new biological mechanisms are resolved, these can be converted into molecular pathways and associated with KEs for which that was not known so far. Second, by introducing more specific ontological annotations of AOP-Wiki content, new literature can be discovered, allowing a dynamic process of describing the biology behind the AOP. Furthermore, by improving annotations of AOP content and making those annotations available through RDF [58], the automatic creation of molecular AOPs based on AOP-Wiki contents will be possible. Validation of the method is required by comparing it with other transcriptomic-based analysis methods or other *in vitro* methods that measure KE activation. This could be done through more extensive case studies on well-established AOPs in the AOP-Wiki.

